# Association of avian biodiversity and West Nile Virus circulation in *Culex* mosquitoes in Emilia-Romagna, Italy

**DOI:** 10.1101/2025.08.18.670801

**Authors:** Yiran Wang, Mattia Calzolari, Gianpiero Calvi, Victoria M Cox, Paola Angelini, Michele Dottori, William Wint, Sally Jahn, Giovanni Marini, Ilaria Dorigatti

## Abstract

West Nile Virus (WNV) is a zoonotic arbovirus maintained in a transmission cycle between *Culex* mosquitoes and birds, occasionally spilling over into humans. The impact of avian biodiversity on WNV circulation remains debated, with studies reporting both negative and positive correlations (dilution and amplification effects respectively) across different settings. In Europe, this relationship remains largely unexplored, particularly in regions with high WNV transmission, such as Emilia-Romagna in Northern Italy.

We explored the association between avian biodiversity and WNV circulation in *Culex* mosquitoes in Emilia-Romagna using 11 consecutive years (2013–2023) of entomological surveillance data paired with two avian data sources. We calculated avian biodiversity indices (Shannon’s, Simpson’s, and Chao2) from observation records from the Farmland Bird Index project and applied linear regression models to assess their relationship with WNV circulation levels. Moreover, we used Bayesian spatiotemporal regression models and gridded weekly avian abundance estimates from the eBird project, to analyse the associations between avian species richness indices and WNV-positivity rates at 68 geolocated mosquito traps across the region.

We observed consistent negative associations between WNV circulation in the *Culex* population and avian biodiversity indices, supporting the dilution effect hypothesis (DEH). We found that non-passerine species richness was negatively associated with WNV mosquito circulation, whereas passerine species richness exhibited a positive association after adjusting for environmental factors and spatial random effects. These findings suggest that passerines may amplify WNV transmission, while the presence of non-passerine species is associated with reductions in WNV circulation.

This study provides the first empirical evidence supporting the DEH for WNV in Europe. These findings have important implications for avian surveillance and future WNV modelling studies, and can inform regional, national and European policies on biodiversity conservation and public health planning.

**Author summary:** West Nile Virus (WNV) is a zoonotic, vector-borne disease maintained in a mosquito-bird-mosquito transmission cycle. The role of avian biodiversity in WNV transmission remains debated, with studies reporting both dilution effects (negative associations) and amplification effects (positive associations) across different settings. Here, we examined the impact of avian biodiversity on WNV circulation in *Culex* mosquitoes in Emilia-Romagna, Italy, one of Europe’s WNV hotspots. We analysed 11 years (2013–2023) of entomological surveillance data alongside two ornithological datasets, respectively collected by the Farmland Bird Index project and the citizen science project eBird. These different datasets allowed us to test the dilution effect hypothesis (DEH) at broad and fine spatiotemporal resolutions. Our findings provide support for the DEH for WNV, revealing that higher bird community diversity is associated with reduced WNV circulation. Additionally, we found that non-passerine species richness is linked to lower levels of WNV circulation in the vector, whereas passerine species richness is positively associated with WNV transmission from entomological surveillance. These results point towards the potential amplifying role of passerine species and the protective role of non-passerine species in WNV transmission dynamics, which has important implications for species conservation, public health policy and future modelling studies.

## Introduction

West Nile Virus (WNV) is a single-stranded, positive-sense RNA virus of the genus *Flavivirus* (family *Flaviviridae*), first isolated in Uganda in 1937 [1, 2]. It is a zoonotic arbovirus primarily transmitted by *Culex* mosquitoes and maintained through a mosquito-bird-mosquito transmission cycle, with birds serving as main reservoir hosts and humans and horses as dead-end hosts [3]. The relevance of a bird species in WNV transmission primarily depends on the rate of contact with mosquito vectors and its reservoir competence. The rate of contact is largely reflected in mosquito feeding patterns observed in the field, shaped by a combination of mosquitoes’ intrinsic host preference and ecological factors such as host abundance, behaviour, and host-vector interactions [4, 5]. Blood-meal analyses revealed that *Culex* mosquitoes predominantly feed on *Passeriformes* birds [6, 7]. In North America, American Robin (*Turdus migratorius*), House Sparrow (*Passer domesticus*), and House Finch (*Haemorhous mexicanus*), all passerine species, are the most common blood-meal sources [6, 8, 9]. In Europe, common Blackbirds (*Turdus merula*), considered ecologically analogous to American Robins, and House Sparrows are the primary targets [10–12], with Magpies (*Pica pica*) also frequently fed upon in Italy [13]. Reservoir competence refers to a host’s ability to infect feeding mosquitoes and is determined by the magnitude and duration of infectious viremia following WNV infection. Laboratory studies measuring viremia profiles after experimental infections identified *Passeriformes*, particularly from the *Corvidae*, *Fringillidae*, and *Passeridae* families as highly competent WNV hosts [14]. Consequently, passerine birds are generally regarded as the optimal hosts for WNV transmission.

While WNV human infections are mostly asymptomatic (∼80%) or cause mild, self-limiting flu-like symptoms such as headaches and fever (∼20%) , less than 1% of the WNV infections progress to severe West Nile neuroinvasive disease (WNND) which can be fatal, primarily affecting older individuals [15]. Currently, no targeted therapeutics or vaccines are available for WNV infection in humans. In recent years, WNV has caused several outbreaks in Europe [16–18], including a large epidemic in 2018, in which more human cases than the combined total of the previous seven years were reported [16], with a significant number of WNND cases and fatalities. While several modelling studies have investigated the role of environmental drivers and climatic anomalies, such as high spring temperatures, in driving WNV transmission [19–21], less progress has been made on the investigation of the role of biodiversity on WNV transmission, mainly due to difficulties associated with the availability of bird species occurrence and abundance data linked with WNV entomological surveillance data. This knowledge gap has hindered the understanding of WNV disease ecology, which is essential for developing effective prevention and control strategies that align with the One Health framework.

Biodiversity is defined as the natural variety and variability among living organisms, the ecological complexes they inhabit, and their interactions with one another and with the physical environment [22]. It is typically characterized by (i) the number of different entities, e.g., species richness; (ii) the relative abundance of different entities, e.g., species evenness; (iii) the specific identities of different entities, e.g., community composition [23]. Increasing evidence suggests that biodiversity might play a pivotal role in influencing the transmission of infectious diseases, particularly vector-borne zoonoses [24].

In 2000, Ostfeld and Keesing first proposed the dilution effect hypothesis (DEH), which suggests that higher biodiversity within a host community is associated with reduced infection rates in vectors and ultimately lowers the risk of pathogen transmission to humans [24]. This phenomenon may occur through various mechanisms, including a reduction in contact rates that facilitate pathogen transmission, regulation of competent host populations, decreased vector density, and enhanced recovery of infected host individuals [25]. Over the past two decades, the DEH has been demonstrated in multiple vector-borne zoonotic disease systems, such as Lyme disease [25, 26], Tick-borne Encephalitis [27], and, more recently, Usutu Virus [28].

In the case of WNV, there is no consensus on the role of avian host biodiversity in transmission, with findings supporting both dilution and amplification effects. Several studies in the United States support the dilution effect, suggesting that greater avian biodiversity is associated with reduced WNV transmission risk [29–32]. In contrast, more recent studies in North America and Europe have identified amplification effects, where greater avian biodiversity is linked to increased WNV circulation [33, 34]. These conflicting findings and the limited number of European studies underscore the need to investigate the location-specific relationship between avian biodiversity and WNV transmission, which is crucial for understanding the local WNV epidemiology and informing public health practices and policy decisions. Primarily due to the difficulty in obtaining reliable data on the bird community, such analyses remain largely underexplored, including in high WNV transmission regions of Europe, such as Emilia-Romagna in Northern Italy [35].

In this study, we investigate the effect of avian biodiversity on WNV circulation in *Culex* mosquitoes in Emilia-Romagna, Italy, by analysing 11 years of entomological surveillance data from 2013 to 2023 and two ornithological data sources: local bird census data from the Farmland Bird Index project and weekly species abundance estimates from the eBird project. We test the effect of avian biodiversity on WNV circulation as measured by alternative metrics at different spatial and temporal resolutions, adjusting for meteorological and land-use confounders. The insights generated by the analyses presented in this study have the potential to inform bird surveillance programs and guide the integration of biodiversity conservation into public health strategies, both within Emilia-Romagna and in broader contexts.

## Results

### Farmland Bird Index data analysis

Among the 86 mosquito traps included in this study, including 68 traps that were active throughout all surveillance seasons from 2013 to 2023 and 9 traps that were relocated to nearby locations during the study period, 55 traps had matched bird observation records conducted across different times. These traps yielded a total of 659 bird observation records, encompassing 107 species and 26,608 individuals. Of the 107 species, 44 were passerines, accounting for 15,264 individuals. Among the 55 mosquito traps with associated bird observation records, WNV was detected in all traps in at least one year and up to a maximum of ten years. Across these 55 traps, 448 out of 20,981 tested mosquito pools were positive for WNV, 447 of which consisted of *Culex pipiens* specimens, with a single positive pool of *Culex modestus* mosquitoes.

In the Exclusion-based Rarefaction approach, where WNV circulation groups with fewer than 20 bird observations were excluded and sample-based rarefaction was performed (Fig 1b), linear regression results indicated that both rarefied Shannon’s and Simpson’s diversity indices were negatively associated with the number of years in which mosquito surveillance traps detected at least one WNV-positive pool (Shannon: β = ―8.89, p = 0.005; Simpson: β = ―59.04, p = 0.006) (Figs 2a and 2b). This suggested that higher diversity in the bird community was associated with lower WNV circulation in *Culex* mosquitoes. While a negative trend was also observed between bird species richness (Chao2 index) and WNV circulation, this relationship was not statistically significant (β = ―0.15, p = 0.376) (Fig 2c).

**Fig 1.**
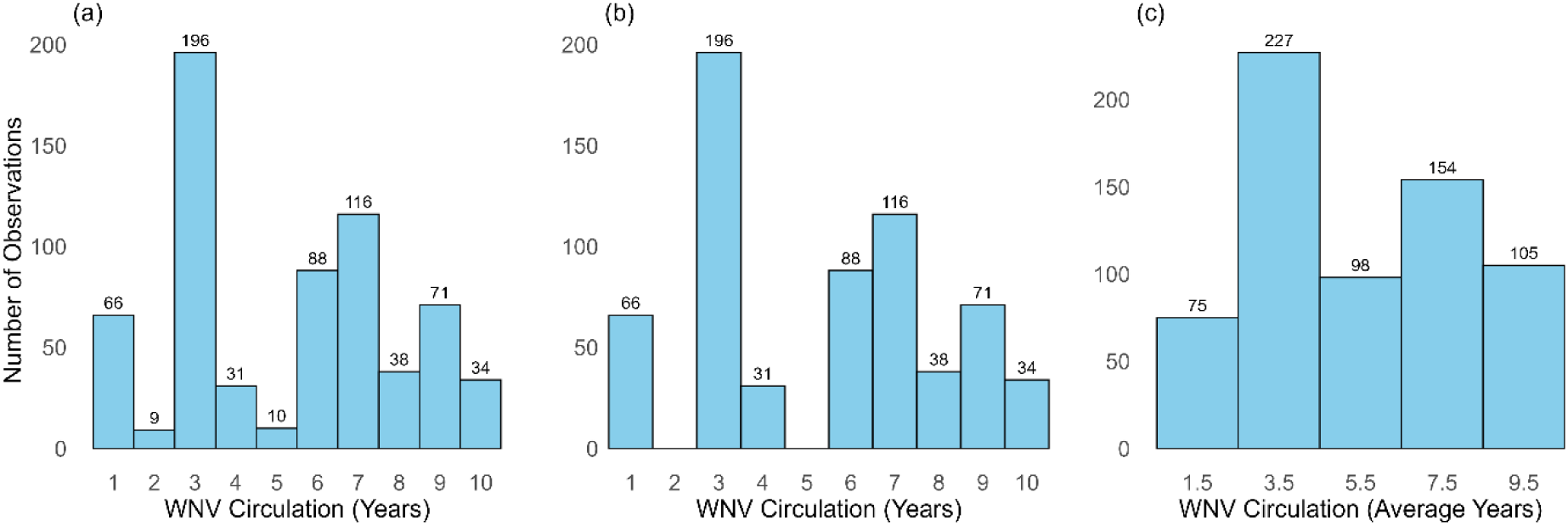
Number of bird observations categorized by WNV circulation groups, based on the number of years with at least one WNV-positive mosquito pool. (a) Classification using the original group definitions. (b) Classification after excluding groups with fewer than 20 observations. (c) Classification with the original two consecutive groups (as shown in panel a) combined.

**Fig 2.**
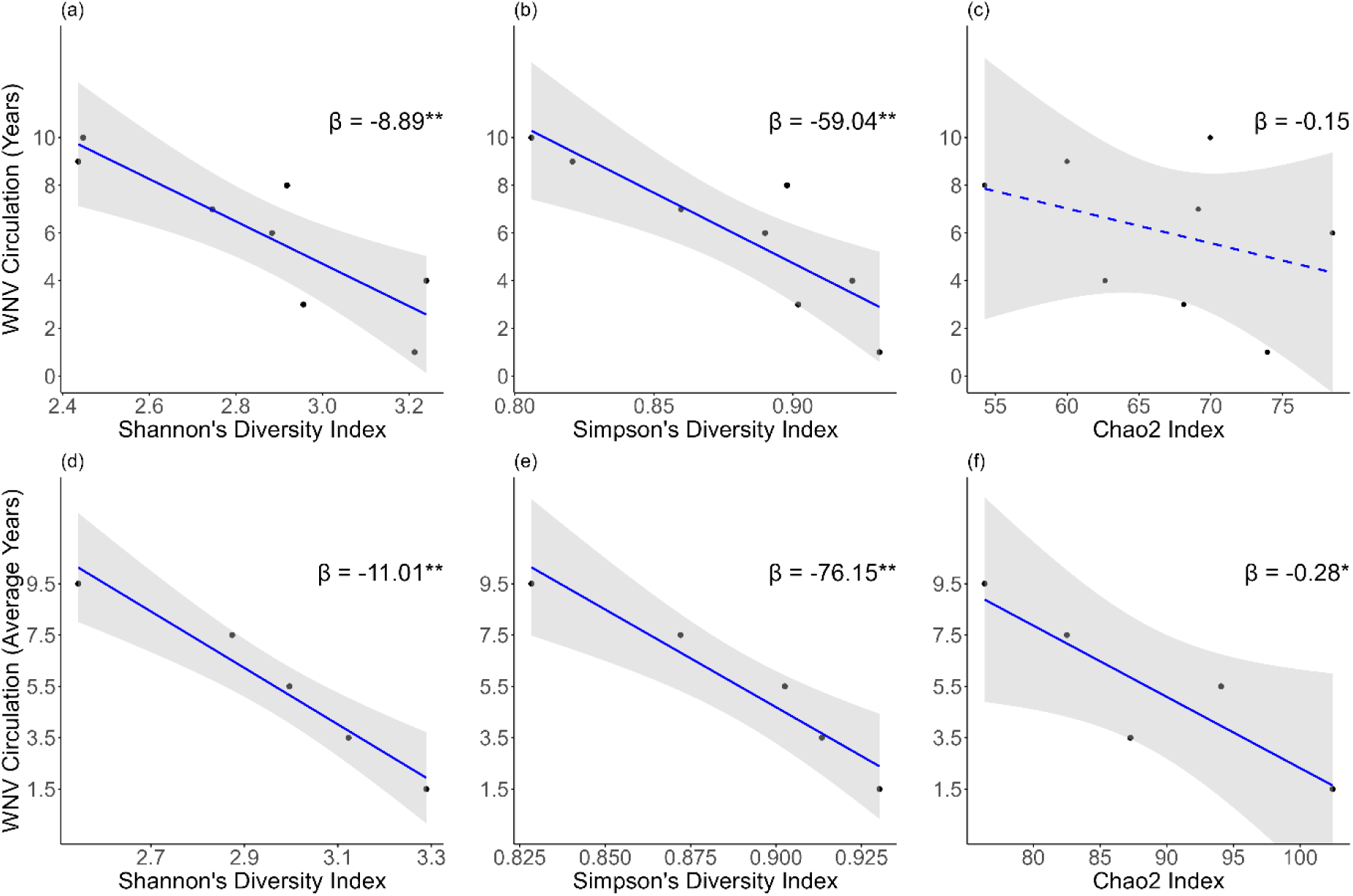
Associations between rarefied bird community biodiversity indices and WNV circulation, measured as the number of years with at least one WNV-positive pool in the Exclusion-based Rarefaction approach (a-c) and the average number of years with at least one WNV-positive pool in the Combination-based Rarefaction approach (d-f). *β* represents the coefficient from simple linear regression, with * indicating a p-value (*p*) *p* ≤ 0.05 and ** indicating *p* ≤ 0.01. Solid lines were used for *p* ≤ 0.05, a dashed line denotes a non-significant trend (*p* > 0.05). (a) Rarefied Shannon’s Diversity Index versus WNV circulation (Exclusion-based Rarefaction). (b) Rarefied Simpson’s Diversity Index versus WNV circulation (Exclusion-based Rarefaction). (c) Rarefied Chao2 Index versus WNV circulation (Exclusion-based Rarefaction). (d) Rarefied Shannon’s Diversity Index versus WNV circulation (Combination-based Rarefaction). (e) Rarefied Simpson’s Diversity Index versus WNV circulation (Combination-based Rarefaction). (f) Rarefied Chao2 Index versus WNV circulation (Combination-based Rarefaction).

In the Combination-based Rarefaction approach, consecutive WNV circulation groups were combined and sample-based rarefaction was performed based on 75 subsamples (Fig 1c).

Similarly to the Exclusion-based Rarefaction method, the results from simple linear regressions revealed negative associations between Shannon’s and Simpson’s diversity indices and WNV circulation (Shannon: β = ―11.01, p = 0.004; Simpson: β = ―76.15, p = 0.008) (Figs 2d and 2e). Additionally, a negative relationship between the Chao2 index and the average number of years with at least one WNV-positive pool was observed, in this case significant (β = ―0.28, p = 0.044), suggesting that higher species richness in the bird community was associated with lower WNV circulation in vectors (Fig 2f). The rarefied biodiversity indices (Shannon’s, Simpson’s, and Chao2) derived from the Exclusion-based and Combination-based Rarefaction approaches are presented in S1 and S2 Tables, respectively.

A sensitivity analysis, conducted using data only from the mosquito traps that were active throughout the entire study period (i.e., excluding the 9 traps that were relocated), showed consistent results across both rarefaction methods (see S1 Text).

### eBird data analysis

The weekly species richness from May to September at 68 mosquito surveillance traps, active throughout all years, derived from the eBird weekly abundance estimates of 148 bird species in the Emilia-Romagna region, exhibits clear spatiotemporal heterogeneity (Fig 3). Overall, species richness is higher in correspondence with the migration periods, when several non-breeding species stopover in the study area for refuelling. At the beginning of the study period in May, the pre-breeding migration is at its final stage, which translates into a slight decline of species richness. From June to July, richness remains relatively stable, coinciding with the end of the breeding season, and then increases again between August and September, due to postnuptial migration. Throughout the season, non-passerine species consistently outnumber passerine species around the mosquito surveillance traps. Among the 45 bird species classified as fully migratory (see S3 Table), only 12 were passerines.

**Fig 3.**
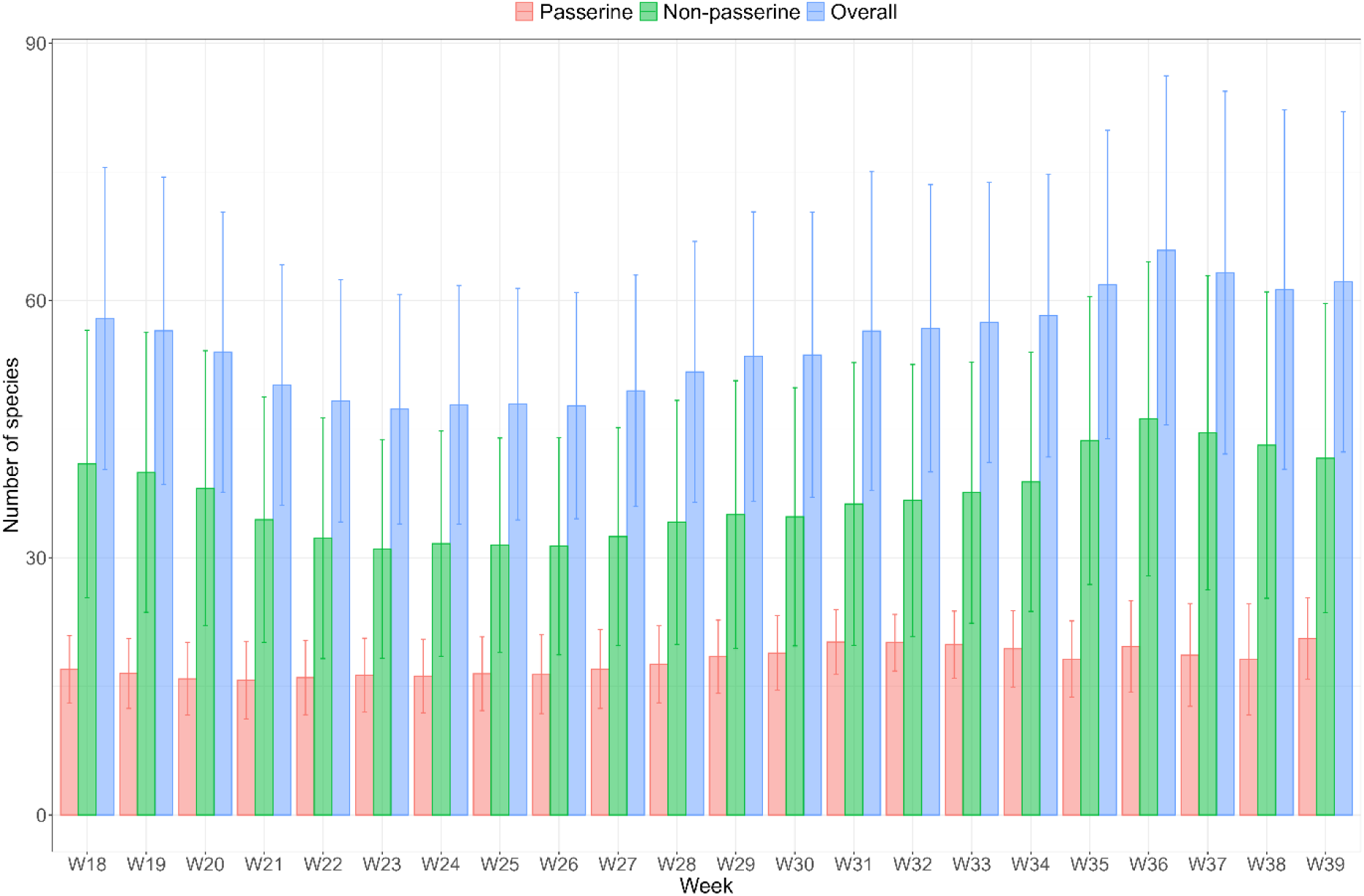
Seasonal trends in weekly passerine (red), non-passerine (green), and overall bird species richness (blue) around 68 active mosquito surveillance traps in Emilia-Romagna, Italy (weeks 18–39, May to September), obtained from eBird abundance estimates. The bars represent the mean values, with error bars indicating the 95% confidence intervals.

Variables included in the spatiotemporal regression model and their associated estimated coefficients are presented in Table 1. The results indicate that non-passerine species richness was negatively associated with the number of mosquito pools testing positive for WNV, after adjusting for the number of pools tested, other covariates, and the spatial random effect. This suggested that a greater presence of non-passerine species within the bird community was linked to reduced WNV circulation in *Culex* mosquitoes. In contrast, passerine species richness was positively associated with the number of WNV-positive pools, indicating that a higher presence of passerine species was correlated with increased WNV circulation in the vector population. Additionally, the proportion of artificial surfaces was negatively associated with WNV circulation, while the weekly average temperature at a three-week lag and cumulative precipitation of the current week were positively associated. To assess model fit, the mean absolute error (MAE) was calculated for both the final model and a baseline model containing only the intercept and spatial random effect. The MAE for the final model was 0.496, compared to 0.899 for the baseline model, indicating that the inclusion of species richness variables alongside environmental (presence of artificial surfaces) and meteorological (temperature and precipitation) fixed effects substantially improved model performance.

**Table 1.** Summary of fixed effects from Bayesian spatiotemporal regression: posterior mean and 95% credible intervals (CrI) for predictor variables.

| Variables | Mean | 2.5% CrI | 97.5% CrI |
| --- | --- | --- | --- |
| Non-passerine species richness (lag 0) | -0.034 | -0.049 | -0.020 |
| Passerine species richness (lag 0) | 0.098 | 0.054 | 0.143 |
| CLC1 (Artificial surfaces) | -4.147 | -6.023 | -2.270 |
| Weekly Average Temperature (lag 3) | 0.516 | 0.448 | 0.584 |
| Weekly Cumulative Precipitation (lag 0) | 0.018 | 0.000 | 0.036 |

Further analysis focusing on migratory birds revealed that the species richness of fully migratory birds, with a two-week lag, was significantly negatively associated with the number of WNV-positive mosquito pools, after adjusting for temperature, artificial land use, and the spatial random effect (S4 Table).

## Discussion

Our findings support the DEH of WNV transmission in Emilia-Romagna, Italy, and points toward the potential role of passerine birds in enhancing WNV circulation and non-passerine birds in mitigating its transmission.

In the Farmland Bird Index data analysis, we found that Shannon’s and Simpson’s diversity indices were consistently significantly lower around mosquito surveillance traps with higher WNV circulation levels, as classified by the number of years with at least one WNV-positive mosquito pool from 2013 to 2023 in Emilia-Romagna. These two indices reflect the relative proportions of different species in the community, with lower values indicating simpler communities dominated by fewer species. This suggests that more diversified bird communities, with more complex trophic and ecosystem interactions, may be less permissive to WNV circulation, while simplified communities are likely more prone to viral transmission, which is consistent with findings from studies conducted in the USA [30–32]. For instance, negative associations were observed between Shannon’s diversity index (calculated from 1998 breeding bird survey data) and human WNV incidence in both 1998 and 2002 in the eastern USA [30]. Similarly, another study found that bird diversity, measured by Shannon’s index, was negatively associated with mosquito WNV prevalence in the Saint Louis, Missouri region and with national human WNV incidence in the early 2000s [31]. Furthermore, an analysis of 20 years of North American Breeding Bird Survey (BBS) data (1989–2008) reported a negative correlation between Shannon’s index and mortality in American crow, a species highly susceptible to WNV, with prior studies indicating severe regional population declines that coincided with the spread of the virus [32]. In addition, we observed a significant negative association between bird species richness, estimated via the Chao2 index, and WNV circulation in *Culex* mosquitoes when using the Combination-based Rarefaction method. The results obtained on the analysis of the eBird data are consistent with previous findings from Louisiana, USA, where non-passerine richness was also negatively associated with WNV incidence in *Culex* mosquitoes [29]. Together, these findings corroborate the DEH of WNV transmission in Emilia-Romagna.

There are several potential mechanisms underlying the DEH in vector-borne disease systems, which could operate in concert in nature including: (i) transmission reduction (or encounter reduction), which occurs when the presence of additional species lowers the rates of contact between vectors and the species that facilitate pathogen transmission; (ii) susceptible host regulation, referring to scenarios where the presence of more species limit or regulate the number of highly susceptible or competent hosts, often through interspecific competition for limited resources; (iii) vector regulation, defined as a decrease in vector density due to the presence of more bird species feeding on mosquitoes, which could cause higher mosquito mortality or other constraints on the vector population; and (iv) recovery augmentation, which occurs when the presence of additional species enhance the recovery of infected individuals by providing resources, such as acting as mutualists or prey [36]. Our findings align most closely with the encounter reduction theory, supported by the strong negative associations between Shannon’s and Simpson’s diversity indexes and WNV circulation intensity observed in the Farmland Bird Index analysis, as well as the significant negative relationship between the species richness of fully migratory bird species and the number of WNV-positive mosquito pools in the eBird analysis. These results suggest that the introduction of additional bird species, possibly through migration and resulting in increased diversity of the bird community, may lower WNV circulation in *Culex* mosquitoes in our study site. Notably, Emilia-Romagna hosts a higher proportion of non-passerine species than passerine species during the WNV transmission season, as indicated by both the bird observations from the Farmland Bird Index Project and the eBird data. In addition, most of the fully migratory species identified in the region are also non-passerines.

Additionally, our study provides statistical evidence linking greater passerine species richness to increased WNV circulation, which may imply a potential amplifying role of passerine birds in the spread of WNV. This finding is consistent with a study conducted in Southwest Spain, where a higher number of passerine species in the bird community was associated with increased prevalence of Usutu virus, a pathogen that can co-circulate with WNV and is also primarily transmitted by *Culex* mosquitoes [37]. This pattern may be explained by both higher reservoir competence of passerine species and/or their more frequent contact with mosquito vectors. While laboratory studies have demonstrated that many passerine species develop high viremia levels following WNV infection, these findings are largely based on North American species and the NY99 virus strain prevalent in the USA [38], which may not be generalizable to other WNV strains or geographic contexts [39]. In Europe, a few studies have assessed the reservoir competence of local bird species using European WNV strains, and there is some but inconsistent evidence that European passerines exhibit high viremia levels as those observed in studies from North America [40–42]. In contrast, a study analysing 150 European passerine species found that models incorporating life-history traits affecting their rate of contact with mosquitoes, such as body mass, nest height, and life stage, better explained variations in bird seroprevalence than models based solely on species identity. This suggests that in Europe, variations in vector-host contact rates might play a vital role at shaping WNV transmission [43]. Therefore, the positive association observed between passerine species richness and WNV circulation in our study may reflect the frequent feeding of *Culex* mosquitoes on passerine birds, as supported by previous blood meal analyses conducted in both North America and Europe [6, 8–13]. Nevertheless, further targeted studies are needed to better characterise the reservoir competence of European passerines and clarify their role in WNV transmission.

In terms of environmental factors, the proportion of artificial surfaces was found to be negatively associated, whereas weekly average temperature was positively associated with mosquito WNV circulation; both patterns are consistent with the literature [44–46]. A weak positive association was also observed between weekly cumulative precipitation and WNV presence in *Culex* mosquitoes, which may be explained by the influence of precipitation and adverse weather conditions on migratory bird stopover behaviour, thereby potentially altering the composition of bird communities during migration periods [47].

Compared with previous studies which focused on single transmission seasons [29–31, 34], this study analysed 11 years of mosquito surveillance and bird observation data. Additionally, we performed sample-based rarefaction to adjust for differences in sampling effort across trap locations and used the Chao2 index to estimate species richness, which accounts for unobserved rare species due to sampling limitations, providing a more accurate measure than simple species counts [29, 30, 34]. Moreover, in the eBird data analysis, we were able to examine the role of bird species richness in modulating the spatiotemporal dynamics of WNV circulation while comprehensively accounting for the confounding effects of environmental factors, which is novel and possible due to the resolution of the eBird data.

A key limitation of the Farmland Bird Index data analysis is that it was conducted using aggregated trap data rather than individual trap-level observation records. While this ensured sufficient bird observations to derive biodiversity indices at each WNV circulation level, it limited the extent to which we could account for potential confounders such as meteorological and land-use factors at fine spatial resolution. A potential drawback of the eBird data is that abundance estimates are based on the eBird citizen science project which by design does not systematically survey pre-defined locations and hence may introduce some bias. Furthermore, these estimates reflect seasonal trends and do not capture interannual variations. Additionally, because eBird abundance estimates are relative and standardised within each species, they do not allow for direct comparisons between species, limiting our analysis to species richness rather than more comprehensive biodiversity metrics. As a result, while our findings show a positive association between passerine species richness and WNV circulation in *Culex* mosquitoes, it remains challenging to disentangle whether this reflects a genuine amplifying role or simply their higher availability as blood-meal sources, considering that *Passeriformes* is the largest avian order. This limitation is not unique to eBird data but is common to most bird data sources, given the difficulty of accounting for heterogeneity in detection and reporting rates across species.

To date, the only study investigating the DEH in the case of WNV transmission in Europe, conducted in southern Spain in 2013, found a positive association between avian species richness and WNV seroprevalence in house sparrows [34], a known competent reservoir host [40]. The authors attributed this to *Culex perexiguus*, the primary WNV vector in the region, being more abundant in rural areas where avian biodiversity tends to be higher compared to urban settings. In our study site, the primary vector is *Culex pipiens*, and our analysis is based on virus presence in mosquito vectors rather than WNV seroprevalence in birds. This difference may explain the divergent findings, on top of regional variations in ecological context and host–vector interactions that can further contribute to the contrasting results.

The analyses presented in this study support the DEH for WNV transmission in Emilia-Romagna, providing the first such evidence in a European setting. These insights could inform ornithological surveillance efforts by guiding enhanced monitoring in areas characterized by high passerine abundance or low avian biodiversity. Furthermore, our findings will inform the development of mechanistic models of WNV transmission, most of which currently consider general bird compartments or only competent bird hosts [48, 49]. In addition, the analytical approach developed in this study highlights the value of generating high-resolution bird diversity and abundance maps, which can help improve our understanding of biodiversity’s role in disease dynamics and point towards a beneficial role of avian biodiversity conservation to reduce the local long-term risk of WNV circulation in the bird, equids, and human populations.

Finally, our study underscores the importance of coordination between disease surveillance and bird monitoring programs to facilitate the collection of targeted data for understanding the circulation patterns of WNV and other zoonotic diseases. Future studies can build on the results presented in this study to further explore the mechanisms underlying these associations and to assess how biodiversity conservation efforts may influence the environmental circulation of WNV.

## Conclusions

In this study, we explored the relationship between avian biodiversity and WNV circulation in *Culex* mosquitoes in Emilia-Romagna, Italy by analysing 11 years of mosquito surveillance data (2013–2023) together with data from the Farmland Bird Index project and the eBird project. We found not only negative associations between bird community diversity, as measured by Shannon’s and Simpson’s diversity indices, and WNV circulation in the vector community, but also that non-passerine species richness was linked to reduced WNV circulation, while passerine species richness was associated with increased circulation. These findings provide evidence of the DEH for WNV in Europe and suggest the potential amplifying role of passerine species and the protective effect of non-passerine species in WNV transmission. These results have implications for ornithological surveillance, future modelling studies, and for the development of One Health public health policies considering the role of biodiversity conservation on human and animal health.

## Materials and Methods

### Study area

Emilia-Romagna region covers around 22,450 km² and has a resident population of about 4.47 million people [50]. From 2009, Emilia-Romagna adopted the national surveillance arbovirus plan locally and implemented an enhanced integrated surveillance program targeting mosquitoes, animals and humans which has been continuously refined over time and is still active to date [50, 51]. Surveillance activities focus on an approximately 11,000 km² area located in the Po Valley plain, where over 90% of the region’s residents live. This area is characterized by optimal ecological conditions for WNV circulation [50].

### Mosquito surveillance data

Mosquito surveillance in Emilia-Romagna was initiated after the first human WNV case was reported in the region in 2008 [52] and since then has been conducted annually during the summer months (June–October). Initially, in 2009, mosquito collections were performed at both fixed and temporary stations, with sampling frequencies ranging from weekly to monthly. From 2010 onward, the collection process was standardized, with fixed geo-referenced stations sampled fortnightly [50]. The surveillance network comprehensively covers the regional plain area using a grid of 11 km × 11 km cells, with traps strategically located to optimize mosquito collection. Female mosquitoes were captured using carbon dioxide (CO₂) baited traps or gravid traps [50]. Since August 2017, only CO₂ baited traps have been employed. Each trap is activated for a single night per collection, after which captured mosquitoes are counted, identified to species, and pooled based on date, location, and species, with a maximum of 200 individuals per pool. The pooled samples are subsequently tested for WNV presence using real-time polymerase chain reaction (RT-PCR) [50].

### Farmland Bird Index data analysis

#### Bird observation data

The bird observation data used in this analysis were collated by Lipu (Italian BirdLife partner) within the Farmland Bird Index project funded by the Ministry of Agriculture, Food Sovereignty and Forestry through the European Agricultural Fund for Rural Development [53, 54]. Data collection is based on a stratified sampling design with single-visit point counts [54]. Italy was divided into 10 km× 10 km squares (UTM projection), and within each square, point counts were conducted in 15 of the 100 1 km × 1 km cells, which serve as the sampling units for this study. Observations were carried out using unlimited-distance point counts [55], where trained observers recorded bird species and abundance through both visual and auditory detections, regardless of the distance from the observation point, in a 10-minute count period. These surveys were conducted annually between the beginning of May and the end of June (beginning of July in mountain regions) during morning hours [54]. Further methodological details of the project are available on the website: www.reterurale.it/farmlandbirdindex.

#### Mosquito and bird data integration

68 mosquito traps that were active during all surveillance seasons from 2013 to 2023 were included in this study. The intensity of WNV circulation at each trap was quantified as the number of years with at least one mosquito pool testing positive for WNV, based on the mosquito surveillance data. Bird observation records from the same period were extracted from sampling units located within a 3-km buffer of the selected mosquito traps (Fig 4). This buffer was chosen to align with the flight range of *Culex* mosquitoes [56].

**Fig 4.**
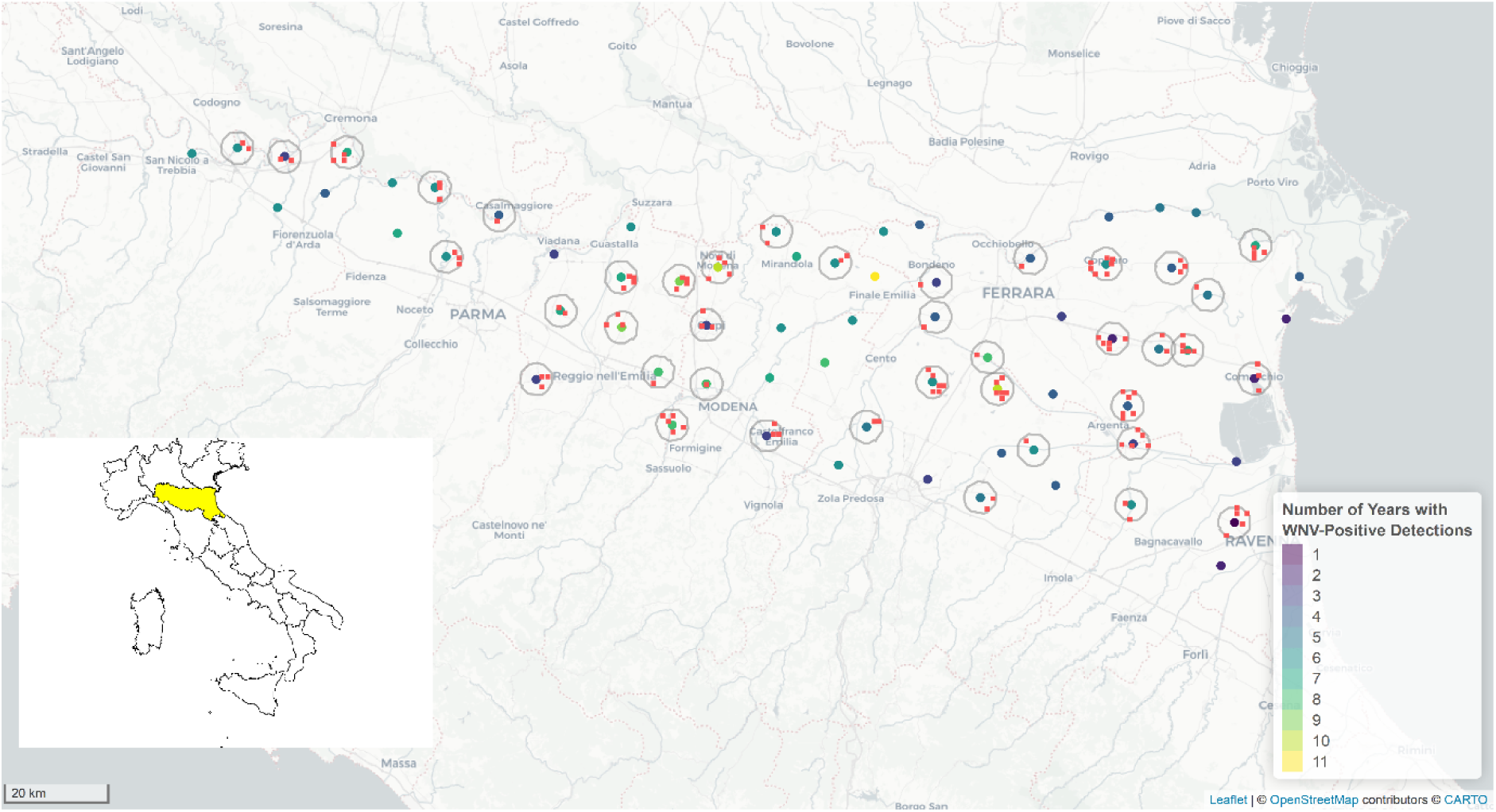
Map of WNV mosquito surveillance traps active from 2013 to 2023 (points, color-coded from purple to yellow by WNV circulation level, classified by the number of years with at least one mosquito pool testing WNV positive) in Emilia-Romagna (highlighted yellow on the map of Italy), with corresponding buffer areas (grey circles) and bird observational stations (red squares). Some traps do not have associated buffer areas because no matched bird observation data were available.

#### Group classification and rarefaction

Mosquito traps were grouped according to the number of years in which at least one WNV-positive mosquito pool was detected. Bird observation records within the buffer zones surrounding these traps were then compiled for each group. In the main analysis, we included 9 mosquito traps that were relocated to nearby sites within the same surveillance grid during the study period alongside the 68 traps that were active throughout the entire study period. A sensitivity analysis was conducted using only data from the 68 active traps (see S1 Text).

To account for differences in the number of observations across groups when calculating biodiversity indicators, sample-based rarefaction was applied [57, 58]. This method involves repeatedly drawing *n* samples without replacement from each assemblage (i.e., set of samples), where *n* is the smallest number of samples among all assemblages [58]. The mean biodiversity indices calculated across these repeated subsamples represent the expected values for each assemblage. In this analysis, rarefaction was performed with 1,000 iterations.

The number of observations across WNV circulation group is shown in Fig 1a and minimum threshold for reliable sample-based rarefaction was considered to be 20 [59]. In the Exclusion-Based Rarefaction approach, rarefaction was performed with a minimum of 31 observations across the remaining groups (Fig 1b). In the Combination-Based Rarefaction approach, consecutive groups were combined, and rarefaction was conducted using 75 subsamples (Fig 1c).

#### Biodiversity indicators

For each group, three rarefied biodiversity indicators were calculated:

1. Shannon’s Diversity Index (*H*) [60]:

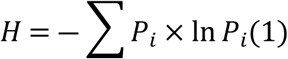 Where *P*_*i*_ is the relative abundance of species *i*. A larger value of *H* suggests higher diversity.
2. Simpson’s Diversity index (1 ― D) [61]:

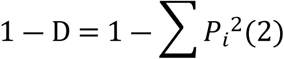 Where *P*_*i*_is the proportional abundance of species *i*. D represents the probability that two randomly selected individuals belong to the same species. A higher value of 1 ― D indicates greater diversity.
3. Chao2 Index (*Ŝ*_*Chao*2_) [62]:

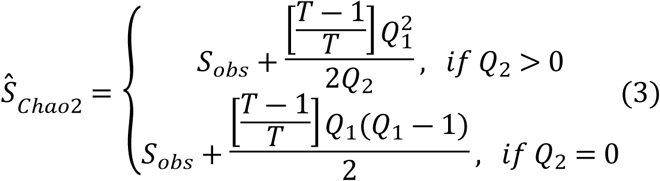 Where S_obs_ is number of observed species, T is number of samples, Q_1_ is number of species observed only once, and Q_2_ is number of species observed only twice. The Chao2 index is a non-parametric species richness estimator accounting for unseen species based on the frequency of rare ones, which is particularly useful in situations where not all species are observed due to sampling limitations [62].

The Shannon and Simpson diversity indices were calculated using the *vegan* R package [63].

#### Statistical analysis

Simple linear regressions were performed to explore the relationships between the three biodiversity indicators and WNV circulation intensity in *Culex* mosquitoes.

### eBird data analysis

#### eBird data

This analysis used data from the eBird status and trends project developed by Cornell Lab of Ornithology [64, 65], which applies machine learning models to estimate distributions, relative abundance, and population trends at high spatial and temporal resolutions across the full annual cycle for over 10,000 bird species. These estimates are generated by analysing the relationships between bird observations from eBird, a large-scale citizen science project, and habitat variations while accounting for biases and noise inherent in community science data, such as variability in bird observer behaviour and effort.

Weekly relative abundance data with a 3 km × 3 km spatial resolution for May to September (calendar weeks 18–39) were downloaded from the eBird website (https://science.ebird.org/en/status-and-trends) for bird species classified within the Emilia-Romagna region. eBird provides two versions of estimates: the 2021 version (based on data from 2007–2021) for some species and the 2022 version (based on data from 2008–2022) for others. Given that the population distribution of one species is largely consistent across years and only the presence/absence status is of interest, we downloaded data for species with no missing values during the study period, which reflect seasonal trends in the region.

Of the 308 bird species identified in eBird within the Emilia-Romagna region, 148 had complete relative abundance data during the study period and were included in the analysis (see S3 Table). The 148 species reported in eBird encompassed 90% of the top 30 species reported in the Farmland Bird Index project within a 10 km buffer of the mosquito surveillance traps from 2013 to 2023, and 80% of the top 50 species ranked by detection frequency. Weekly abundance estimates for the 148 species were transformed into binary presence/absence data and aggregated into weekly species richness values, including overall bird species richness as well as passerine and non-passerine richness, as illustrated in Fig 3. In a sensitivity analysis, we explored the role of migratory birds in WNV circulation using the same analytical framework described above and weekly species richness estimates calculated exclusively for species classified as fully migratory in the study region.

#### Environmental and meteorological data

Meteorological data, including daily measurements of mean, minimum, and maximum temperature, precipitation, relative humidity, solar radiation, leaf wetness hours, and evapotranspiration, were obtained from the ERG5 dataset provided by the Regional Agency for Prevention, Environment, and Energy of Emilia-Romagna, Italy (ARPAE) [66, 67] for the entire study period from 2013 to 2023. For each meteorological variable, weekly averages or cumulative values were computed and then averaged over the study period to obtain seasonal trends.

Land use proportions around mosquito traps were derived from the 2012 Corine Land Cover (CLC) dataset provided by the European Environment Agency, which classifies land cover into five categories: artificial surfaces (CLC1), agricultural areas (CLC2), forests and semi-natural areas (CLC3), wetlands (CLC4), and water bodies (CLC5) [68]. For each trap location, land use types within a 3 km buffer were identified by extracting CLC data from shapefiles provided by the Institute for Environmental Protection and Research (ISPRA) using QGIS (www.qgis.org). The proportion of each land cover class within each mosquito trap’s buffer zone was then calculated.

#### Spatiotemporal analysis

A Bayesian spatiotemporal model was developed to examine the relationships between WNV circulation in *Culex* mosquitoes and bird species richness, adjusting for environmental covariates in a seasonal context using the Integrated Nested Laplace Approximation - Stochastic Partial Differential Equation (INLA-SPDE) approach implemented in R-INLA (www.r-inla.org) [69, 70]. The model assumes that mosquito WNV circulation is a continuous spatially-correlated process across the region, and represents this using a discretized Gaussian Markov random field (GMRF) based on a triangulated mesh [71, 72].

The model combines the spatial field (a random effect) with the avian, meteorological, and land use variables (fixed effect covariates) to reconstruct the observed patterns in mosquito WNV circulation. Details on the mesh and prior distributions for the GMRF parameters are provided in S2 Text. Analyses were conducted using the *R-INLA* package (version 24.05.10).

For the 68 mosquito traps that were active throughout each surveillance season and consistently sampled during calendar weeks 18 to 39 from 2013 to 2023, the total number of mosquito pools tested for WNV and the number of pools testing positive for WNV were aggregated by weeks across all years from 2013 to 2023. The weekly count of WNV-positive pools was taken as the response variable, with the total number of pools tested adjusted accordingly. A total of 87 initial variables were included in the analysis, including the bird species richness (overall, passerine, non-passerine), and meteorological and land use covariates. Species richness and meteorological variables were extracted with up to 3-week lags to account for lag effects.

To identify key predictor variables, we employed Spike and Slab, a Bayesian variable selection method [73] performed using the *spikeslab* R package [74]. Pairwise correlations among the selected variables were then calculated, and highly correlated variables were excluded to mitigate multicollinearity. For pairs of variables within the same category (e.g., temperature-related variables) with a correlation coefficient exceeding 0.6, the variable with the higher Bayesian model average (BMA) value was retained. For pairs of variables from different categories (e.g., temperature and precipitation), priority was given to removing compound variables (e.g., evapotranspiration).

Following this procedure, five variables (*X*_{*i*,*j*},*t*_, j=1, …, 5)— namely, passerine species richness at zero-lag, non-passerine species richness at zero-lag, weekly average temperature at a three-week lag, weekly cumulative precipitation at zero-lag, and land use type CLC1 (artificial surfaces) —were included as fixed effects in the final spatiotemporal model.

Although CLC2 (agricultural areas) was initially retained alongside these five variables after the variable selection process, it was not statistically significant and was therefore excluded from the final model to avoid potential bias from including a non-informative covariate.

Given the high proportion of zero observations (70.11%) in the response variable, we applied a zero-inflated binomial distribution to account for the two types of zeros: structural zeros (representing locations where no WNV-positive pools should occur) and sampling zeros (representing locations where no WNV-positive pools were detected due to sampling limitations). The model was specified as follows:

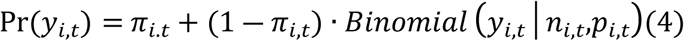

where *y*_*i*,*t*_is the number of WNV-positive mosquito pools observed at trap *i* in week *t*; *π*_*i*,*t*_represents the probability of a structural zero; *n*_*i*,*t*_ is the number of pools tested at trap *i* in week *t*; and *p*_*i*,*t*_is the predicted probability of a pool testing positive, which was defined as shown in eq. 5:

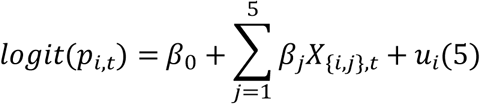

where *β*_0_is the intercept; *β*_*j*_is the regression coefficient for the predictor *X*_{*i*,*j*},*t*_at trap location *i* and week *t*; and *u*_*i*_is the spatial random effect at trap location *i*. All analyses were performed using R version 4.4.0 [75]. The data and code used in this analysis are provided in: https://github.com/yiranwangic/WNV_biodiversity_EM.

## Acknowledgments

We are grateful to all the observers involved in the bird census.

## Supporting information

**S1 Table. Rarefied biodiversity indices (Shannon’s, Simpson’s, and Chao2) for eight WNV circulation groups based on years with at least one WNV-positive mosquito pool (Exclusion-based Rarefaction).**

**S2 Table. Rarefied biodiversity indices (Shannon’s, Simpson’s, and Chao2) for five WNV circulation groups based on average number of years with at least one WNV-positive mosquito pool (Combination-based Rarefaction).**

**S3 Table. Bird species in Emilia-Romagna, Italy with available weekly abundance estimates from May to September to download from the eBird website.** The entry ‘21’ in column ‘Version’ indicates the 2021 version of eBird (based on data from 2007–2021) and ‘22’ indicates the 2022 version (based on data from 2008–2022). The species classified as fully migratory are highlighted in bold.

**S4 Table. Summary of fixed effects from Bayesian spatiotemporal regression examining the association between fully migratory bird species richness and West Nile Virus (WNV) circulation in Culex mosquitoes: posterior means and 95% credible intervals (CrIs) for predictor variables.**

**S1 Text. Sensitivity analysis for Farmland Bird Index data.**

**S2 Text. Mesh construction and prior specifications of the INLA-SPDE model.**

